# Metagenomic insights into the metabolism and ecologic functions of the widespread DPANN archaea from deep-sea hydrothermal vents

**DOI:** 10.1101/2020.02.12.946848

**Authors:** Ruining Cai, Jing Zhang, Rui Liu, Chaomin Sun

**Author notes:** Corresponding author: Chaomin Sun.

## Abstract

Due to the particularity of metabolism and the importance of ecological roles, the archaea living in deep-sea hydrothermal system always attract great attention. Included, the DPANN superphylum archaea, which are massive radiation of organisms, distribute widely in hydrothermal environment, but their metabolism and ecology remain largely unknown. In this study, we assembled 20 DPANN genomes comprised in 43 reconstructed genomes from deep-sea hydrothermal sediments, presenting high abundance in the archaea kingdom. Phylogenetic analysis shows 6 phyla comprising Aenigmarchaeota, Diapherotrites, Nanoarchaeota, Pacearchaeota, Woesearchaeota and a new candidate phylum designated DPANN-HV-2 are included in the 20 DPANN archaeal members, indicating their wide diversity in this extreme environment. Metabolic analysis presents their metabolic deficiencies because of their reduced genome size, such as gluconeogenesis, *de novo* nucleotide and amino acid synthesis. However, they possess alternative and economical strategies to fill this gap. Furthermore, they were detected to have multiple capacities of assimilating carbon dioxide, nitrogen and sulfur compounds, suggesting their potentially important ecologic roles in the hydrothermal system.

**IMPORTANCE:** DPANN archaea show high distribution in the hydrothermal system. However, they possess small genome size and some incomplete biological process. Exploring their metabolism is helpful to know how such small lives adapt to this special environment and what ecological roles they play. It was ever rarely noticed and reported. Therefore, in this study, we provide some genomic information about that and find their various abilities and potential ecological roles. Understanding their lifestyles is helpful for further cultivating, exploring deep-sea dark matters and revealing microbial biogeochemical cycles in this extreme environment.

## INTRODUCTION

Archaea are regarded as a significant part in the microorganism, playing key roles in the energy flow and biogeochemical cycle (1–3). Originally, all archaea were found by cultivation and assigned to two clades: the Crenarchaeota and the Euryarchaeota (4). With novel sequencing techniques like metagenome generating, the archaeal tree have been dramatically expanded (5). To date, at least four supergroups are contained in archaea kingdom: Euryarchaeota, TACK, Asgard, and DPANN archaea (4). Not only that, archaeal genes and functions are greatly uncovered for better understanding their evolutions, lifestyles and ecological significances (6). Among the archaeal domain, the DPANN are a superphylum firstly proposed in 2013 (7). They consist of at least 10 phylum-level lineages discovered: Altiarchaeales, Diapherotrites, Aenigmarchaeota, Pacearchaeota, Woesearchaeota, Micrarchaeota, Parvarchaeota, Nanoholoarchaeota, Nanoarchaeota and Huberarchaea, and are the remarkable aspects of the life tree (5, 8–10). Recently, the lifestyles and ecology of the DPANN group collected from different biotopes were analyzed and summarized through culture-independent methods (7, 8, 11, 12).

Firstly, in the metabolism researches, their primary biosynthetic pathways including amino acids, nucleotides, lipids and vitamins were detected to be absent (10). Furthermore, they mostly lack complete tricarboxylic acid cycle (TCA cycle) and pentose phosphate pathway, suggesting they are deficient in the ATP production (8). But they have been hypothesized to utilize a putative ferredoxin-dependent complex 1-like oxidoreductase as an alternative pathway (11). Similarly, genes encoding enzymes involved in the generation of fermentation products including lactate, formate, ethanol and acetate were found in many DPANN genomes, strongly suggesting that they might have evolved to use substrate-level phosphorylation as a mode of energy conservation (8, 12). Secondly, due to their small genomes and limited metabolism, many members of DPANN were inferred to be dependent on symbiotic interactions with other organisms and may even include novel parasites (8). Several reports recognized that the Nanoarchaeota and the Parvarchaeota required contacting with similarly or larger sized hosts to proliferate or parasitize (13–16). The Huberarchaea were also described as a possible epibiotic symbiont of Candidatus Altiarchaeum sp. (17). Thirdly, their ecologic significance was only studied by a few reports and limited in the Woesearchaeota despite their wide existence. The Woesearchaeota were revealed a potential syntrophic relationship with methanogens and deemed to impact methanogenesis in inland ecosystems (7). Moreover, Woesearchaeota were also found to be ubiquitous present in the petroleum reservoirs, where they contribute to the ecosystem diversity and the biogeochemical cycle of carbon (18).

As mentioned above, the insights of the DPANN group collected from different biotopes have been studied, such as groundwater (19), surface water (20) and marine sediments (21). However, the DPANN living in deep-sea hydrothermal vent sediments were little researched, even though showing high abundance (22–24). Deep-sea hydrothermal vent sediments provide the extensive habitats for microbial lives, and these microbes also drive nutrient cycling in this ecosystem (25, 26). Furthermore, the microflora living there are presumed or detected to have specific capabilities because of their special habitats (27). Most significantly, they are considered as crucial hints to understand special life processes and ecosystems in the extreme environments. Given that the DPANN take majority proportion in the archaea kingdom of hydrothermal system, it’s meaningful to know their lifestyles, metabolism even ecologic functions in the global biogeochemical cycles.

Here, we found the DPANN were abundant in the deep-sea hydrothermal vents of the western Pacific and then investigated their carbon cycles, assimilation and dissimilation of nitrogen and genomic features about their special living environments. Finally, the DPANN were detected to have multiple capacities of metabolizing carbon, nitrogen, and sulfur compounds, playing crucial roles in the hydrothermal system, even though having reduced genome and limited amino acids and nucleotides synthesis abilities.

## RESULTS

### The DPANN show high abundance and wide diversity in the hydrothermal system

To explore the composition and specificity of archaeal communities inhabiting in the deep-sea hydrothermal vent sediments, we collected and sequenced three sediment samples from the Okinawa Through in the western Pacific (Fig. S1, Table S1). After assembly, 43 draft genomes were reconstructed using coverage binning. All assembled genomes represent >50% completeness (26 genomes > 70%) and < 10% contamination (DATASET S1).

For classifying these genomes, phylogenetic tree was constructed using 37 single-copy protein-coding marker genes (Table S2). Overall, 43 genomes distribute in 19 lineages basing on the phylogenetic distance analysis (branches length distance < 0.6) (Fig. S2, DATASET S2). Thereinto, 20 DPANN genomes belonging to different phyla were discovered (Fig. 1), showing high abundance in the archaeal kingdom of the hydrothermal vent sediments.

**FIG 1.**
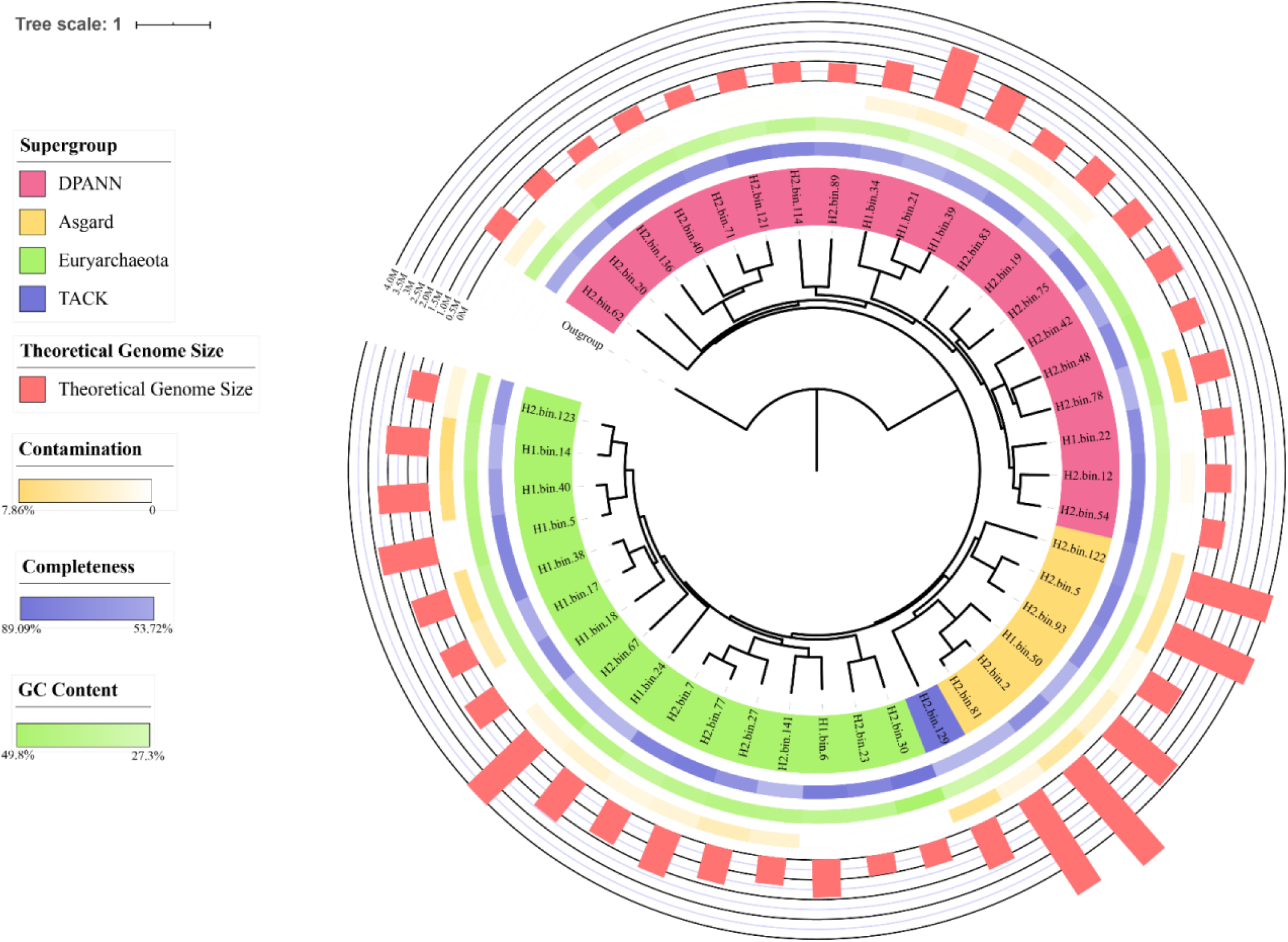
Phylogeny of 43 assembled genomes from hydrothermal vent sediments. The maximum likelihood tree base on 37 single-copy protein-coding genes. Different supergroup is colored in red (DPANN), yellow (Asgard), green (Euryarchaeota) and blue (TACK) within the corresponding leaves in the tree, respectively. Uncolored leave is outgroup. Outer colored circles indicate the completeness (blue), GC content (green), contamination (yellow) and theoretical genome size (red) of assembled genomes, respectively. The reference information is available in DATASET S1.

In order to determine the confirmed phylum of the DPANN group living in hydrothermal vents sediments (named DPANN-HV in this study), phylogenetic analysis was done using the same methods above (Fig. S3, DATASET S3). Totally, the 20 genomes represent in 6 phyla comprising Aenigmarchaeota (n=1), Diapherotrites (n=1), Nanoarchaeota (n=2), Pacearchaeota (n=3), Woesearchaeota (n=9) and a new candidate phylum designated DPANN-HV-2 (n=4). In the tree of the DPANN group, the DPANN-HV-2 form a distinct clade and have no lineage with a closer similarity (Fig. S3).

Further supports for designating DPANN-HV-2 as a new candidate phylum stem from directly comparing the average amino acid identity (AAI) with all public DPANN genomes from NCBI (Fig. 2, DATASET S4). The DPANN-HV-2 are respectively < 46.11% AAI and are more similar to themselves than to other phyla. The AAI are as low as values observed among other disparate phyla in DPANN as well as the threshold considered for separate phyla (28) (Fig. S4, DATASET S4). These evidences further prove that they are a new phylum in the DPANN group suggesting the great specificity of DPANN in such an extreme environment. More importantly, their genome information enables us to speculate their lifestyles and metabolic versatility of these extreme hydrothermal communities especially DPANN group.

**FIG 2.**
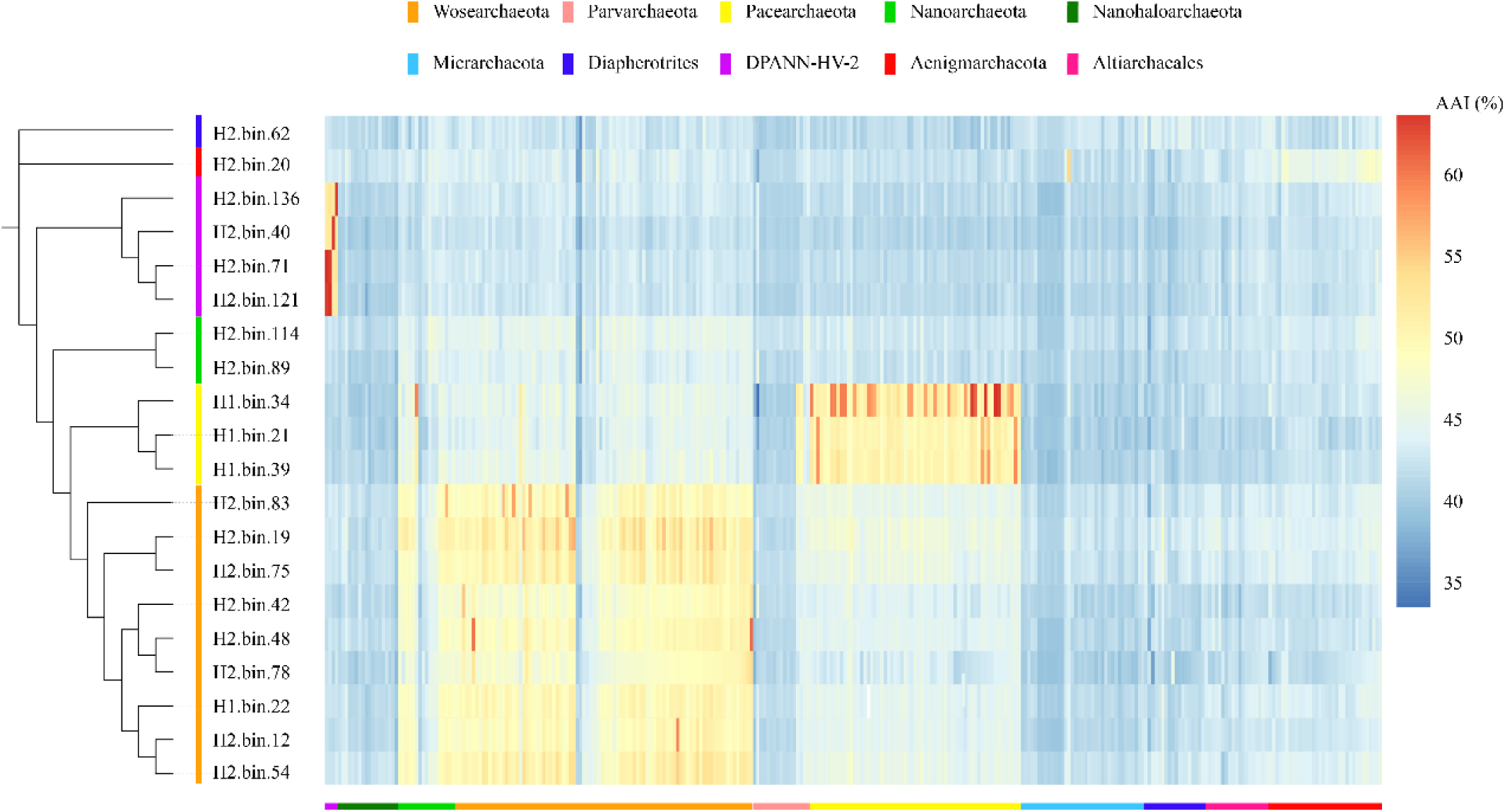
Genome-identity correlation matrix of assembled genomes compared with referenced DPANN genomes. The referenced genomes containing all DPANN genome sequences from NCBI. The amino acid identity correlation matrix was calculated by CompareM. All data are available in DATASET S4.

### The DPANN-HV lack gluconeogenesis pathway but possess alternative carbohydrates metabolizing and carbon fixation capabilities

In the investigation into the lifestyles and metabolisms of the DPANN-HV, we found their genomes were smaller than other archaeal genomes (Fig. S5, DATASET S3). Therefore, we inferred their limited or economical physiological capacities by assigning metabolic functions to proteins in each genome. Firstly, we investigated their abilities to metabolize carbohydrates. Glycolysis pathway plays an essential role in most organisms to offer energy through breaking down glucose. We searched genomes for three glycolysis key enzymes glucokinase (*glk*), phosphofructokinase (*pfk*) and pyruvate kinase (*pyk*)) encoding genes, and found at least one gene in 90% genomes (Fig. 3, DATASET S5). And pyruvate kinases producing ATP in glycolysis are expressed by more than 90% members. It means that the DPANN-HV may be able to utilize a part of glycolysis to obtain energy. Reverse to this pathway, gluconeogenesis, which produces sugars (namely glucose) for catabolic reactions from non-carbohydrate precursors, is also crucial in most organisms. But unexpectedly, 75% members of DPANN-HV lack the pyruvate carboxylase (pc) meaning that they can’t use gluconeogenesis pathway through pyruvate (Fig. 3). Therefore, it proves that gluconeogenesis pathway is absent in the DPANN-HV.

**FIG 3.**
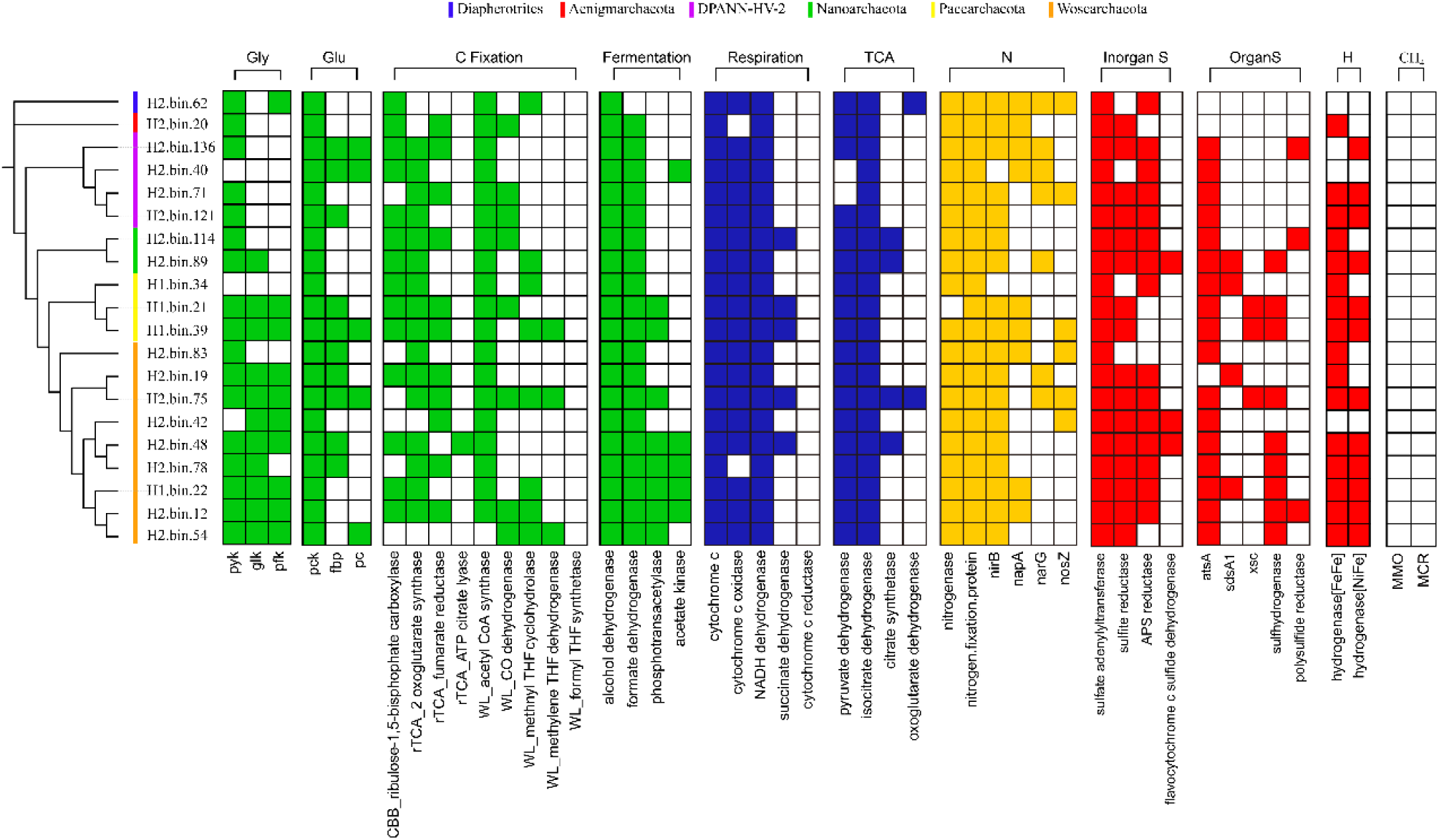
Core metabolic genes detected in DPANN-HV genomes. The presence of core metabolic genes involved in carbon metabolism, respiration process, nitrogen metabolism and sulfur metabolism was investigated. All gene predictions are based on the sequences alignments with KEGG, GenBank NCBI NR and UniProt database. Solid colors: this gene exist in this genome; White: this gene is absent in this genome. Key metabolic predictions and gene abbreviation list are supported by the gene information in DATASET S5.

However, the significance of gluconeogenesis is supplying organisms glucose to produce energy when starving. So how do the DPANN-HV, the archaea lacked of gluconeogenesis, get sugar? To address this question, we investigated the abilities of the community to degrade and metabolize complex carbohydrates, peptides and alkanes. To assess the degradation capacities of complex sugars, the CAZy database was used for searching carbohydrate-active enzymes (CAZymes) in the genomes of DPANN-HV. We found that there were about 70 CAZymes per genome (Fig. 4, DATASET S6). Generally, the DPANN-HV encode glycoside hydrolases and esterases more and polysaccharide lyases less. Only 27 polysaccharide lyases were aligned, which consist of pectin degraded PL1 and unknown PL0, suggesting that the DPANN-HV had limited polysaccharide decomposition functions reacted by polysaccharide lyases. But approximately 12% CAZymes are potentially secreted (Fig. S6), meaning that the complex substrates are broken down outside the cell and taken up later. Theses secreted enzymes including GH13, GH43, GH23, GH18 and so on (Fig. S7), are able to help the DPANN-HV to degrade complex carbohydrates (glucan, xylose etc.) in the surroundings, which can be further used for cell life. Conclusively, the DPANN-HV are able to utilize complex substrates as carbon source by secreted carbohydrates-active enzymes. Additionally, secreted peptidases encoding genes distribute widely in the DPANN-HV genomes (Fig. S8 and S9, DATASET S6) suggesting that some peptidases are secreted to decompose proteins in the environment. Among these peptidases, collagenases serve as breaking down collagen from environments. Accordingly, owing to the abundance of giant tube worms in hydrothermal vents (29), collagen are rich in this system and supply sufficient carbohydrates for the DPANN-HV, which supply enough substrates for collagenase. Additionally, the DPANN-HV are capable of degrading aromatic hydrocarbons (Fig. S10). The enzyme, PPS (phenylphosphate synthase), can be used to transform the phenol to phenylphosphate (30). Then the products are converted to fumaric acid used in respiration process through PPC (phenylphosphate carboxylase) and other enzymes (31). However, the key enzyme, PPC, was not detected in the DPANN-HV genomes, suggesting that there might be any new aromatic hydrocarbon utilization pathways in this group. Given that hydrothermal vent sediments contain rich aromatic hydrocarbons (32), thus, the aromatic hydrocarbon degradation abilities of the DPANN-HV reflect their adaptabilities to hydrothermal environments, and also contribute to the assimilation of deep-sea organic matters and the carbon cycle in the hydrothermal systems.

**FIG 4.**
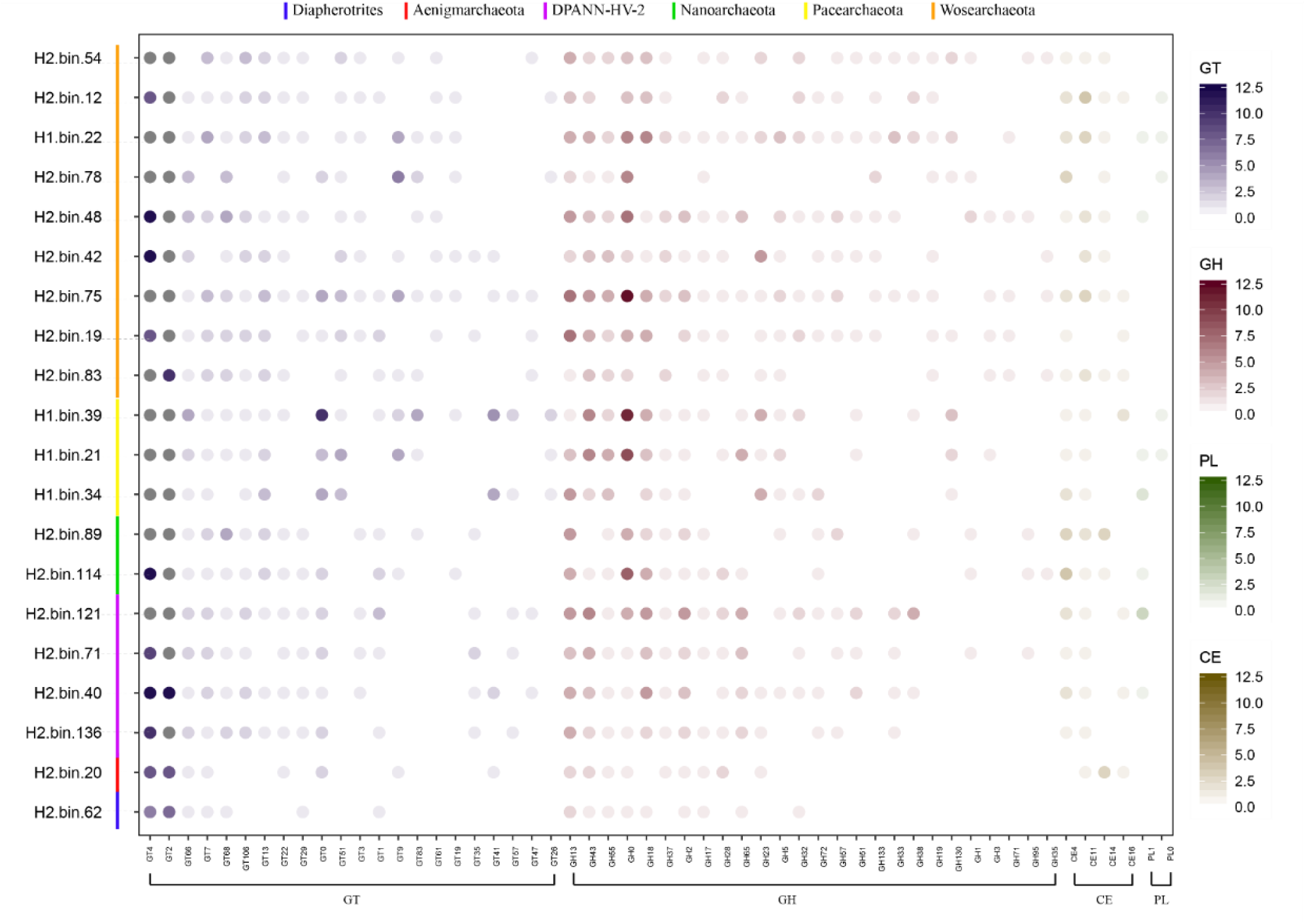
Relative abundance of genes encoding for carbohydrate-active enzymes (CAZymes) encoded in the DPANN-HV genomes. The number of carbohydrate esterases (CE), glycoside hydrolases (GH) and polysaccharide lyases (PL) encoded in the DPANN-HV summarized for each genome was investigated. Numbers of genes belonging to different CAZyme families per genome are presented by colors in circles. The gene distribution and classification are supported by the information in DATASET S6.

Conclusively, we found the DPANN-HV applied several pathways to metabolize extracellular carbon source (carbohydrates, proteins, and alkanes) existing in the environment. But considering the harsh condition in the deep-sea, we sought to ask when the environmental nutriment is poor, how will the DPANN-HV synthesize glucose? Carbon fixation pathways probably are good solutions.

To assess the presence of carbon fixation, we built a gene database containing core enzymes of the Calvin-Benson-Bassham cycle (CBB cycle), the reductive citric acid cycle (rTCA cycle) and the Wood-Ljungdahl pathway (WL pathway). Commonly, the DPANN-HV have CBB cycle and incomplete rTCA cycle (Fig. 3 and 5). And a few genomes were detected to encode genes of WL pathway such as H2.bin.39, H2.bin.54 and H2.bin.75. Even though, these genomes have different pathways, the carbon fixation abilities are undoubted because the carbon dioxide fixation genes are present in all genomes. Moreover, multiple key enzymes for carbon dioxide fixation pathways appear in the same individual, although these pathways are incomplete. We infer that the DPANN-HV adopt multiple pathways to assimilate more carbon sources and use less enzymes as well as less consumption. Therefore, they can convert inorganic carbon to the organic carbon substances efficiently for not only their own growth but other organisms’. Because of their carbon assimilation and high abundance, the DPANN-HV are proposed to play “producer” like roles in lightness deep-sea, and are probably important links in the element cycle and energy transfer of the hydrothermal system.

**FIG 5.**
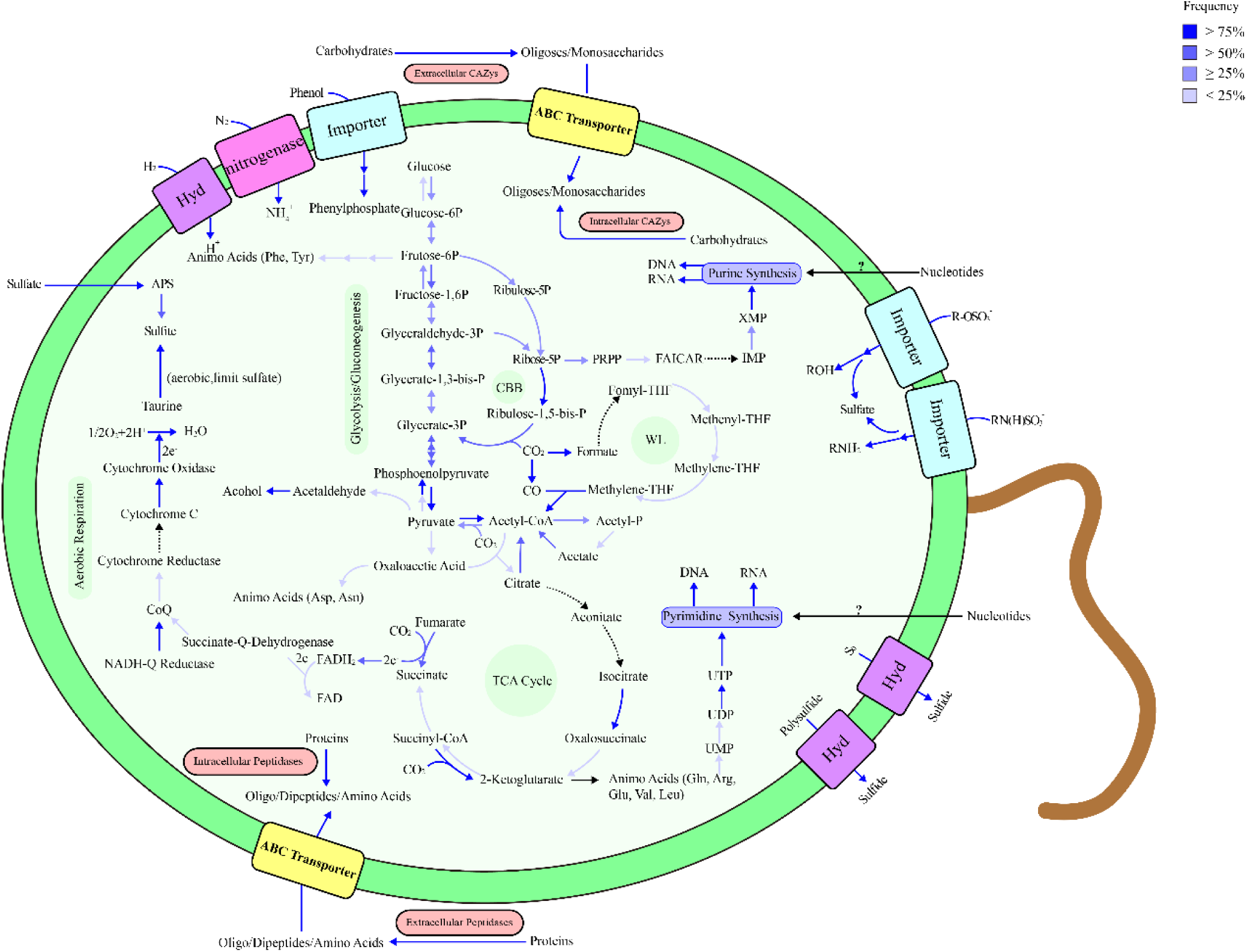
Inferred physiological capabilities of the DPANN-HV. The predictions of metabolic pathways are generated by the KEGG annotations and core genes’ analyses above. The dashed lines indicate missing pathways and the lines with different colors mean the presence frequency of pathways in the DPANN-HV genomes. Details about genes are provided in DATASET S5. Hyd: hydrogenases; APS: adenosine phosphosulfate; WL: Wood-Ljungdahl pathway; CBB: Calvin-Benson-Bassham cycle; THF: tetrahydrofuran; FAICAR: 5-Aminoimidazole-4-carboxamide ribonucleotide; PRPP: Phosphoribosyl pyrophosphate.

### The DPANN-HV assimilate nitrogen but limit in *de novo* nucleotides and amino acids syntheses

Next, we investigated the DPANN-HV for their involvement in nitrogen cycle. The results showed that the nitrogenases distributed in the genomes of DPANN-HV (Fig. 3). In addition, some necessary genes encoding regulators for the nitrogen fixation system were also found in most genomes, suggesting that the DPANN-HV were able to fix nitrogen and produce ammonium salts. Ammonium salts, which are regarded as one of most important nutrients, might not only satisfy with the need of the DPANN-HV but also other lives’ in the same environment. Such a phylum what execute biological nitrogen fixation process must be essential in keeping balance of community structure in the ecosystem.

The main purposes of compounds containing nitrogen in the livings are synthesizing nucleotides and amino acids. However, such a group with nitrogen fixation commonly lack of both complete pathways. Through the analysis of purine synthesis pathway, the DAPNN-HV don’t have the capacities of the *de novo* purine synthesis (Fig. S11, DATASET S5). But there is no lack of genes of mutual transformations among nucleoside monophosphates, nucleoside diphosphates and nucleoside triphosphates. In the analysis of pyrimidine synthesis, the same result was displayed (Fig. S12). We presume the DPANN-HV achieve some semi-finished products like monophosphates from surrounding and then process them to DNA/RNA. If this deduction is the fact, we propose that the DPANN-HV prefer to live in high biomass areas like hydrothermal vents (32), because they need to achieve semi-finished nucleotides from other livings. As for amino acids synthesis, after detecting the genomes, we found the DPANN-HV had limited genes related to amino acid synthesis (Fig. 5, DATASET S5). Totally, they can’t generate independently 11 basic amino acids comprising isoleucine, histidine etc. Nevertheless, the widespread secreted peptidases existing in DPANN-HV might help them to discompose protein into amino acids transported into cells for protein synthesis.

Logically, the both metabolism patterns, reduced nucleotides and amino acids synthesis, are suitable for the microorganisms including DPANN with small genomes because of low consumption and limited genes. Given that the hydrothermal vents areas always show high biomass and high biodiversity, there is no shortage of nucleotides and amino acids released from organisms. Collectively, free of encoding a large amount of enzymes as well as consuming a lot of energy, getting semi-products from environments is a better and more economical method for the DPANN-HV whose genomes are small.

### The DPANN-HV metabolize sulfur compounds and hydrogen which are rich in hydrothermal ecosystem

Because of the high abundance of archaea kingdom in hydrothermal vent sediments, it is reasonable to propose that the DPANN-HV have some special characteristics about hydrothermal vents. It was reported that methane, hydrogen, and compounds containing sulfur were rich in hydrothermal vents (33). We thus searched for the existence of the pathways related to these substances.

To assess whether methane metabolism existed, we aligned genomes to the database consisting of monomethane oxygenase (reacting in aerobic condition) and Methyl coenzyme M reductase (reacting in anaerobic condition) sequences. However, we didn’t find any enzyme about methane metabolism in the DPANN-HV, which indicates that the DPANN-HV might not be methanotrophic microbes. To explore abilities of the DPANN-HV to metabolize hydrogen, hydrogenases in their genomes were aligned, classified, and summarized to their classes, oxygen tolerance and functions. After alignment and classification, only [FeFe] and [NiFe] family hydrogenases were detected but [Fe] family hydrogenases were not (Fig. 3). The latter ones are regarded as methane metabolizing enzymes, their absence also lends credence to the deduction that this superphylum cannot use methane. On the facet of oxygen tolerance, the most of hydrogenases are oxygen-liable, and a few of them are oxygen-tolerant (Fig. S13, DATASET S6). That reflects the DPANN-HV are facultative anaerobes. On the anaerobic condition, hydrogenases work as electron donors and react in fermentation of formate, acetate and alcohol. On the aerobic condition, oxygen-tolerant hydrogenases have the abilities to metabolize polysulfide and convert elemental sulfur or polysulfide into sulfide, which strongly suggests that DPANN-HV might be contributors in the sulfur cycle in the hydrothermal environment.

To reveal the possible roles of DPANN-HV in the sulfur cycle in hydrothermal vent system, we searched their genes which used for utilizing sulfur compounds including inorganic and organic sulfur compounds. The results showed that only a few individuals possess sulfide oxidases, even though sulfide scatters widely in the hydrothermal vents (Fig. 3). However, enzymes catabolizing inorganic sulfur compounds exist widely in this group. For example, sulfate adenylytransferase are present in all genomes, which can convert sulfate, ATP and H^+^ into adenosine phosphosulfate (APS) (34). And then APS is dissimilated to sulfites through APS enzyme, releasing AMP, H^+^ and an oxidative electron receptor (35). In terms of organic sulfur metabolism, we found that *atsA* genes (36) were widely scattered. It suggests the DPANN-HV could metabolize organic sulfur compounds to obtain organic molecules and sulfate for synthesizing cofactors and amino acids (37). The arylsulfatases expressed by *atsA* enable DPANN-HV forage on more kinds of organic sulfur containing substrates (38). It means that the DPANN-HV are more tolerant to these compounds harmful for others. Marine polysaccharides, which are regarded as the most complex organic molecules in the ocean, are highly sulfated comparing to land polysaccharides (39). Marine polysaccharides are rich in the deep-sea hydrothermal system because of its high biodiversity (40). Therefore, these two type enzymes also help DPANN-HV lacking polysaccharide lyases utilize the carbon source better.

The presence of these genes explains that the DPANN-HV are capable of metabolizing many kinds of sulfur compounds in the hydrothermal vents, which shows their adaptabilities to this ecosystem. The DAPNN-HV utilize these substrates and release a large amount of sulfate to offer their own and other livings’ growth. They are also pivotal in the supply of inorganic sulfur even sulfur cycle in the local ecosystem.

## DISCUSSION

In this study, we assembled 20 DPANN group genomes from the deep-sea hydrothermal vents, which took majority in the assembled archaeal genomes. Similarly, high abundance of DPANN was also reported in other hydrothermal systems (24). Moreover, not only in the hydrothermal vent sediments, they exist ubiquitously in estuaries (22), seas (23), inland soils (41) even oligotrophic high-altitude lakes (20) and high-temperature petroleum reservoirs (18). In these researches, the authors detected the operational taxonomic units (OTU), revealed the diversity of archaea and studied potential syntrophic interactions among the archaea in various ecosystems. However, little works illustrated why these lives are widespread and how their metabolisms help them live in such diverse environments. Based on our study, we thought the features of the DPANN-HV group, autotrophy and low consumption, could be a part of answers to these two questions.

The DPANN-HV are essential in hydrothermal ecosystems because of their various significant functions in the cycles of carbon, nitrogen and sulfur, even though their genomes are generally smaller than other archaea. Firstly, they are able to assimilate carbon dioxide. The incomplete reductive citric acid cycle (rTCA cycle) and the Calvin-Benson-Bassham cycle (CBB cycle) were detected in the DPANN-HV genomes. Although they are not complete, the key gene catalyzing carbon dioxide assimilation distributes widely, which strongly suggests that they can transform the inorganic carbon compounds to organic ones, then utilize for life processes and release for other organisms. Most significantly, due to their carbon fixation metabolism, they can serve as producers and a crucial part in the biologic chain of the dark ocean. Besides of carbon fixation, they are capable of assimilating nitrogen. Nitrogenases were detected widely in the DPANN-HV genomes, and the other important regulating genes of nitrogen fixation were also found, suggesting the group had nitrogen fixation system. Nitrogen is metabolically useless to most microorganisms. But it can be converted into ammonia or related nitrogenous compounds which are metabolized by most organisms by the DPANN-HV. On account of their high abundance, the DPANN-HV undoubtedly supply a large amount of nitrogen nutrients and make a great contribution to the nitrogen cycle even biogeochemical cycles. Finally, organic sulfur metabolizing genes were found in the DPANN-HV genomes. Sulfur compounds are known to play crucial roles in the sulfur cycle in hydrothermal vents. The DPANN-HV are able to metabolize polysulfide and phenol sulfate, which can’t be utilized by most microbes. Moreover, the DPANN-HV can produce sulfite and sulfide, the substrates used by most hydrothermal microorganisms. Therefore, they are significant participants in the hydrothermal sulfur cycle. Conclusively, they are essential contributors to maintain hydrothermal ecosystem and promote material circulation and energy cascade. Although many researches mentioned their various functions (7, 8, 11), few focused on their ecological significances. Here, we link the metabolism to ecology as references for further researches.

Due to their reduced genomes, the DPANN-HV tend to use economical strategies. Several important metabolizing pathways in the DPANN-HV groups are incomplete, however, the core genes included in the incomplete pathway or the replaceable metabolism could be detected. We don’t deny the possibility that the absence of the complete pathways is a bias of completeness of genomes, but the possibility would be very little that all 20 genomes commonly miss these pathways. For instance, they mostly lack the complete gluconeogenesis and are unable to catalyze non-carbohydrate precursors to sugars (namely glucose). However, more than 1400 carbohydrates-active enzymes were found in all 20 genomes. And 12% of them can be secreted to the surroundings, degrade polysaccharides and help cell uptake these products. In the same way, the DPANN-HV, which are incapable of *de novo* synthesizing amino acids, can retrieve amino acids from environments using secreted peptidases. As several works revealed (7, 8, 11), we detected the absence of *de novo* nucleotides synthesis as well. However, the DPANN-HV group may possess alternative metabolism to uptake DNA from environments. For example, we found CRISPR specific sites in 75% assembled genomes (DATASET S7). So we infer exogenous nucleotides the DPANN-HV processing are probably from virus though CRISPR immune systems. Besides, type IV secretory pathway presents almost all genomes, meaning the DPANN-HV genomes are able to obtain nucleotides through conjunction (42). Significantly, exosomes exist in 19 DPANN-HV genomes. Bearing proteins, lipids, and RNAs, they mediate intercellular communication (43). Meanwhile, what they carrying are likely to apply substrates for DPANN-HV to synthesize nucleotides and proteins. As some researches inferred (7, 8, 11), these metabolic deficiencies indicated a potential syntrophic and/or mutualistic partnership with other organisms, however, few of them revealed how DPANN group get nucleotides.

Conclusively, owing to their extensive distribution, DPANN-HV are considered to have high adaptabilities in the marine hydrothermal systems. In spite of their reduced genome and limited metabolism in nucleotides and amino acids, they harbor alternative even more economical pathways. It also results from their multiple biological functions such as the assimilation and dissimilation of carbon, nitrogen and sulfur compounds. These features not only nourish them and other organisms in the same environment, but enable them to play a potential key role in this biotope.

## MATERIALS AND METHODS

### Sampling and storage

Sediment samples used in this study were collected by *RV KEXUE* from the western Pacific (124º22’22.794’’E, 25º15’48.582’’N, at a depth of approximately 2190.86 m; and 126º53’85.659’’E, 27º47’21.319’’N, at a depth of approximately 961.24 m) in 2018, and stored at −80 °C.

### Metagenomic sequencing, assembly and binning

Total DNA from 20 g sediments from each sample was extracted using the Tianen Bacterial Genomic DNA Extraction Kit following the manufacturer’s protocol. Extracts were treated with DNase-free RNase to eliminate RNA contamination. Then the DNA concentration was measured by Qubit 3.0 fluorimeter. DNA integrity was evaluated by gel electrophoresis and 0.5 μg of each sample was used to prepare libraries. The DNA was sheared into fragments between 50 - 800 bp using Covaris E220 ultrasonicator (Covaris, Brighton, UK). DNA fragments between 150 bp and 250 bp were selected using AMPure XP beads (Agencourt, Beverly, MA, USA) and then were repaired using T4 DNA polymerase (ENZYMATICS, Beverly, MA, USA). These DNA fragments were ligated at both ends to T-tailed adopters and amplified for eight cycles. Finally, the amplification products were subjected to a single-strand circular DNA libraries.

All NGS libraries were sequenced on BGISEQ-500 platform (BGI-Qingdao, China) to obtain 100 bp paired-end raw reads. Quality control was performed by SOAPnuke (v1.5.6) (setting: -l 20 -q 0.2 -n 0.05 -Q 2 -d -c 0 −5 0 −7 1) (44). The clean data was assembled using MEGAHIT (v1.1.3) (setting:--min-count 2 --k-min 33 --k-max 83 --k-step 10) (45). After that, metaBAT2 (46), Maxbin2 (47) and Concoct (48) were used to automatically bin from assemblies. Finally, MetaWRAP (49) was used to purified and arranged generated data to get the final bins. The completeness and containment was calculated by checkM (v1.0.18) (50).

### Annotation

Gene prediction for individual genomes was performed using Glimmer (v 3.02) (51). Sequences were deduplicated using CD-hit (v 4.6.6) (setting: -c 0.95 -aS 0.9 -M 0 -d 0 -g 1) (52). Then the genomes were annotated by searching the predicted genes against KEGG (Release 87.0) (53), NR (20180814), Swissprot (release-2017_07) and EggNOG (2015-10_4.5v) using Diamond (v0.8.23) by default, and the best hits were chosen. The CRISPR specific sites and proteins were found using CRISPRCasFinder (54). Additionally, in order to search for specific metabolic genes, we downloaded several databases including CAZY (55), MEROPS (56), AnHyDeg (57) and HydDB (58) for identifying carbohydrate active enzymes, peptidases, anaerobic hydrocarbon degradation genes and hydrogenases. Then these databases were aligned to assembled genomes using Diamond (v0.9.29) with E-value 10e-5 with sensitive mode. Also, the sequences from NCBI (only archaeal and bacterial non-redundant sequences were selected) and UniProt (only reviewed sequences were selected) were downloaded for further metabolic analysis. Then we constructed the database for alignments using Diamond with E-value 10e-5. Protein localization was determined for CAZymes and peptidases using TMHMM web tool (59) with default parameters. Finally, all figures were completed using R (v3.5.1).

### Phylogenetic analyses

For revealing the phylum composition of assembled genomes in the archaea kingdom, we downloaded three sequences per archaeal phylum from NCBI ref genomes using Aspera (v3.9.8). Then, Phylosift (v1.0.1) (60) was used to extract 37 marker genes (Table S2) containing in genomes with automated setting. The sequences were trimed using TrimAl (version 1.2) (61) using gappyout function. Finally, maximum likelihood tree was inferred using IQ-TREE (v1.6.12) (62)with GTR+F+R10 model (-bb 1000) and showed by iTOL (v5) (63).

For investigating the confirmed phylum of the DPANN-HV genomes, all referenced DPANN genomes were downloaded from NCBI using Aspera. The same methods above were used in the phylogenetic analyses of referenced DPANN genomes and DPANN-HV genomes but with GTR+F+I+G4 model when using IQ-TREE.

For inferring the phylogeny of the DPANN-HV genomes, the same methods above were completed but with GTR+F+I+G4 model when using IQ-TREE. All assembled genomes were used to calculate the average amino acid identity across all referenced DPANN genomes from NCBI using CompareM (v 0.0.23) with aai_wf function (64). Then we depicted the result with a heatmap using R (3.5.1).

## ACKNOWLEDGEMENTS

We greatly acknowledge all the authors who contribute to this study. Thanks for all editors and anonymous reviewers for valuable feedbacks and constructive comments. We are also thankful to the Center for High Performance Computing and System Simulation of Pilot National Laboratory for Marine Science and Technology (Qingdao) and Demin Xu from the University of Science and Technology of China for their supports in data analysis. This work was funded by the Strategic Priority Research Program of the Chinese Academy of Sciences (Grant No. XDA22050301), China Ocean Mineral Resources R&D Association Grant (Grant No. DY135-B2-14), National Key R and D Program of China (Grant No. 2018YFC0310800), the Taishan Young Scholar Program of Shandong Province (tsqn20161051), and Qingdao Innovation Leadership Program (Grant No. 18-1-2-7-zhc) for Chaomin Sun.

## DATA AVAILABILITY

All referenced genomes were retrieved from NCBI database. All core genes were obtained from NCBI non-redundant protein database (filter: the sequences only belonging to bacteria and archaea), UniProt reviewed database, CAZY, MEROPS, HydDB and AnHyDeg databases. The data supporting the conclusions of this article is included within DATASET S1-S7. All assembled sequence data and sample information are available at NCBI under BioProject ID PRJNA593679.

